# Identifying lncRNA-mediated regulatory modules via ChIA-PET network analysis

**DOI:** 10.1101/331256

**Authors:** Denise Thiel, Nataša Djurdjevac Conrad, Ria X Peschutter, Heike Siebert, Annalisa Marsico

## Abstract

**Background:** Although several studies have provided insights into the role of long non-coding RNAs (lncRNAs), the majority of them has unknown function. Recent evidence has shown the importance of both lncRNAs and chromatin interactions in transcriptional regulation. Although network-based methods, mainly exploiting gene-lncRNA co-expression, have been applied to characterize lncRNA of unknown function by means of ‘guilt-by-association’ strategies, no method exists which combines co-expression analysis with 3D chromatin interaction data.

**Results:** To better understand the function of chromatin interactions in the context of lncRNA-mediated gene regulation, we have developed a multi-step graph analysis approach to examine the RNA polymerase II ChIA-PET chromatin interaction network in the K562 human cell line. We have annotated the network with gene and lncRNA coordinates, and chromatin states from the ENCODE project. We used centrality measures, as well as an adaptation of our previously developed Markov State Models (MSM) clustering method, to gain a better understanding of lncRNAs in transcriptional regulation. The novelty of our approach resides into the detection of fuzzy regulatory modules based on network properties and their optimization based on co-expression analysis between genes and gene-lncRNA pairs. This results in our method returning more *bona fide* regulatory modules than other state-of-the art approaches for clustering on graphs.

**Conclusions:** Interestingly, we find that lncRNA network hubs tend to be significantly enriched in disease association, positional conservation and enhancer-like functions. We validated regulatory functions for well known lncRNAs, such as MALAT1 and the enhancer-like lncRNA FALEC. In addition, by investigating the modular structure of bigger components we show that we can propose regulatory functional mechanisms for uncharacterized lncRNAs, such FLJ37453, RP11442N24 B.1 and LINC00910.

## Introduction

LncRNAs, an heterogeneous group of non-coding transcripts longer than 200 nucleotides, are expressed in a time- and tissue-specific fashion and have been shown to regulate cellular processes such as development, imprinting, X-chromosome inactivation, cancer and immunity [1, 2]. The discovery of extensive transcription of these non-coding transcripts provides an important new perspective on the centrality of RNAs in gene regulation [3]. To date, next-generation sequencing data generated by several consortia, such as ENCODE [4] or FANTOM5 [3] leads to an estimate of the number of potential lncRNA transcripts of about 20000. Although only a smaller fraction of such transcripts might be functional, and despite the substantial progress in mapping lncRNAs, the detailed functional mechanisms for most of them remain elusive [2]. The gap in the understanding of the functional roles of the lncRNAs has largely been due to their poor evolutionary conservation, thus restricting orthology-based approaches for extrapolating function, but also to the limited ability of tools to characterize lncRNA interactions with either proteins, DNA and RNA on a large scale. Concomitant with the increasing number of lncRNAs, a number of resources collecting and curating functional information about lncRNAs have been built in recent years [5].

It has been shown, among others, that lncRNAs can regulate the expression either of their neighboring genes in *cis*, or of more distant genes in *trans*. LncRNA may function via binding to RNA Binding Proteins (RBPs), such as chromatin regulators that can bind both RNA and DNA, or by interactions with other nucleic acids [6].

Through the years, it has been demonstrated that RNA is required for proper chromatin structure and recruitment of the chromatin-modifying complexes to DNA [7]. A major category of well-studied functional lncRNAs is those implicated in coordinated gene silencing, either in *cis* (e.g. the lncRNA Xist, involved in X-chromosome inactivation) or in *trans* (e.g. HOTAIR). Both XIST and HOTAIR have been shown to mediate epigenetic mechanisms of gene silencing [7, 8].

Genome-scale mapping of histone modifications and enhancer-binding proteins has helped to identify lncRNAs involved in gene activation. Enhancers are regulatory sequences that can activate gene expression, and their function depends on the interplay between DNA sequences, DNA-binding proteins, and chromatin architecture [9]. In the last 5 years, the functional landscape of enhancers has become more complex with the evidence that active enhancers can transcribe structured lncRNAs. A recent study performed LOF experiments and found 7 of 12 lncRNAs affecting expression of their cognate neighboring genes [10]. More recently, HOTTIP, an enhancer-like lncRNA, has been discovered to directly interact and activate the WDR5 protein [7], a key component of the mixed lineage leukemia-Trx complex. In other cases lncRNAs activate a neighboring lncRNA, e.g., JPX regulates transcriptional activation of XIST on chromosome X [7]. Long non-coding RNAs with activating function may recruit transcriptional activators involved in establishment of chromosome looping between the lncRNA loci and regulated promoters, such as the mediator complex [11].

The architectural landscape of the nucleus has a profound influence on gene regulation. Chromosome conformation capture technologies, such as 3C, Hi-C, 4C, Capture-C and ChIA-PET have revealed elements that are distally located either on the same or separate chromosomes, to be proximal in the three dimensional nucleus [12]. The effect of such contacts, especially when they correspond to enhancer-promoter or promoter-promoter interactions, mediated by PolII or other factors is an area of intense research [12]. While previous studies indicated that enhancer mechanisms involve transcription factor recruitment, there is evidence that enhancer-promoter interactions might be induced by chromatin looping and mediated by enhancer-like RNA [7].

Additional evidence on potential functions of lncRNAs have been obtained from methodologies which rely on expression patterns and “Guilt by Association”: transcripts sharing common expression patterns are expected to be co-regulated or share common pathways. Most of these methods build a coding-non-coding co-expression network, in which a node represents a molecule and an edge an expression correlation. Such a network is used to identify cellular modules involving both protein coding genes and lncRNAs, and the unknown function of lncRNAs is predicted by transferring functional annotation (e.g. GO terms) from protein coding genes [13, 7, 14]. These approaches however do not necessarily reflect direct interactions between lncRNAs and genes, rather statistical associations, and thus do not directly contribute to an understanding of detailed mechanisms of lncRNA-mediated gene regulation.

In this study we focused on lncRNA regulatory functions in the cell nucleus and constructed the chromatin interaction network involving lncRNAs, genes and other genomic regions using ChIA-PET data in the K562 cell line. ChIA-PET (Chromatin Interaction Analysis by Paired-End Tag Sequencing) was introduced in 2009 and combines ChIP with chromatin capture (3C) technology to detect interactions between genomic regions mediated by a transcription factor of interest [15]. Here, we focus on the PolII-mediated chromatin network. A natural representation of theses data amenable to efficient analysis are complex networks, where nodes represent DNA segments or PETs and edges represent ChIA-PET interactions between two PETs. The analysis of chromatin interaction networks has been an area of active research in the last years, but very few studies have employed network analysis and clustering methods to study chromatin interaction networks [12, 16].

For many biological networks, including gene regulatory networks, the evaluation of well-established node characteristics, in particular centrality measures, are highly suitable for identification of functionally essential elements [17]. Similarly, modular organization is believed to be a generic property of such networks, allowing to uncover subnetworks responsible for a specific function. In gene regulatory networks, e.g., modules often correspond to groups of interconnected cis-regulatory elements. To compensate for the size and heterogeneous nature of the chromatin interaction network, we developed a hierarchical analysis approach: starting from the chromatin graph as a whole, we compute centrality properties on the single chromosome level, followed by a focus on the connected components of the chromosome graphs and finally reaching the level of density-based modules, that are amenable to a detailed analysis in their entirety (Fig. 1a). Specifically, to identify these potential lncRNA-mediated functional modules, we implement a modified version of our previously developed Markov State Models (MSM) clustering approach [18, 19], which aims at identifying subgraphs of high connectivity while maximizing the correlation in expression of genes in the same module. This is motivated by the fact that not only network topological properties, but also similarity between expression patterns might help to assign more biologically meaningful putative functions. To our knowledge, this is the first approach that combines the topology of the 3D chromatin network with co-expression analysis to characterize the unknown function of potential regulatory RNAs.

**Figure 1:**
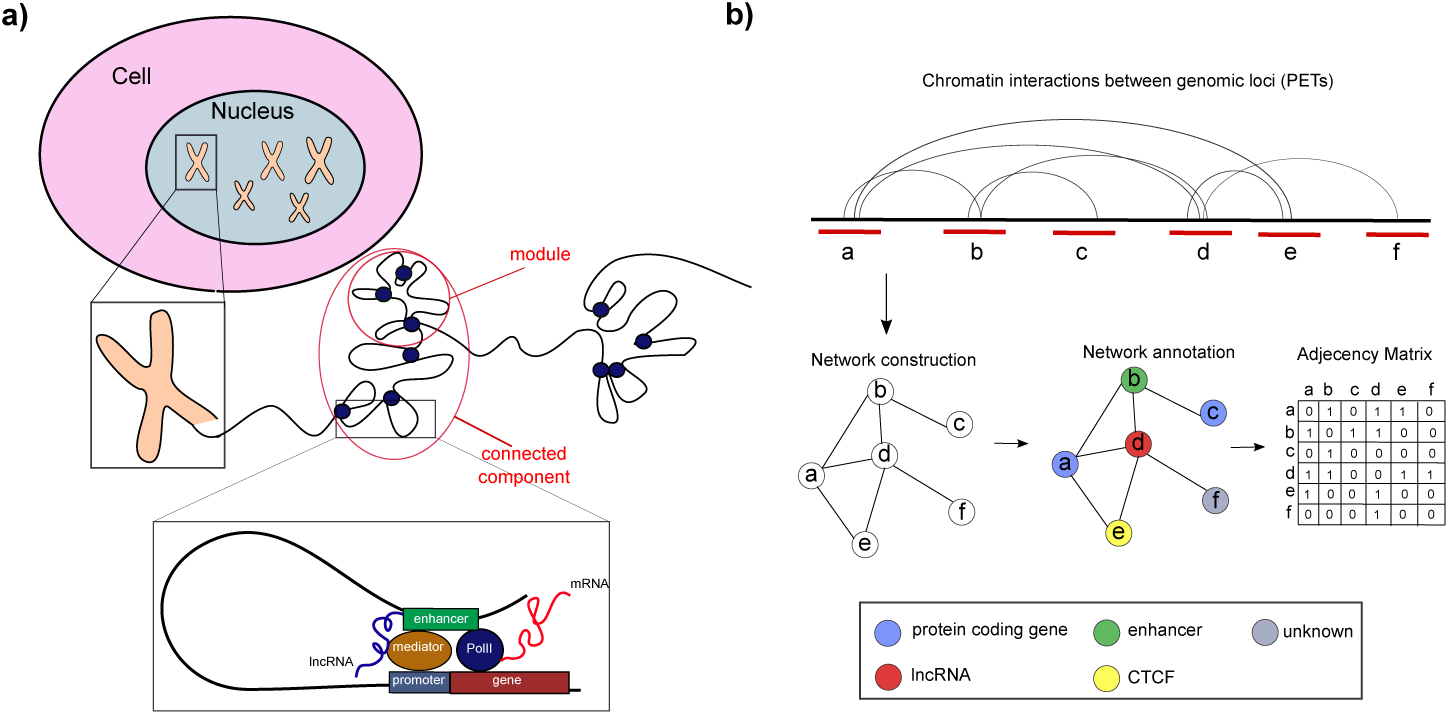
Construction and annotation of the chromatin graph. a) Modular organization of chromatin on each chromosome with highlight on looping between regulatory elements such as enhancers and promoters mediated by PolII, Mediator and nascent lncRNAs. b) Steps involved in network construction and annotation from ChIA-PET data.

We compared our method with other state-of-the-art graph clustering methods, and showed that MSM clustering is superior in returning clusters corresponding to genuine regulatory modules, i.e. whose members exhibit a high correlation in expression between gene-gene and lncRNA-gene. An evaluation of our approach is then conducted by matching distinguished modules and nodes discovered in our analysis with lncRNAs of known importance from the scientific literature, such as ncRNA-a3, FALEC, Xist and MALAT1 [6]. We observe that disease-associated and/or positionally conserved lncRNAs, as well as lncRNAs transcribed from enhancer regions exhibit either high degree or high betweenness centrality, highlighting their crucial role in the leukemia-specific network. Finally, we show the potential of using centrality measures and the module detection for identifying lncRNAs of functional importance in bigger components that cannot be inspected manually. We discuss further potential functional roles for FALEC and MALAT1, as well as attempt to gain functional insights into uncharacterized lncRNAs, such as FLJ37453, RP11442N24 B.1, exhibiting the highest degree in our network, and LINC00910 and LINC00854, involved in CTCF-mediated chromatin looping.

Although some examples are discussed in this manuscript, we propose our approach as general strategy, which can be extended to other cell line-specific ChIA-PET data and to the detailed inspection of the modular structure of other large connected components, as a step forward to annotate lncRNAs with functions in transcriptional gene regulation.

## Methods

### Data collection and Pre-processing

#### ChIA-PET Data

The PolII ChIA-PET interaction network in the K562 cell line was build based on the already processed interaction files downloaded from the ENCODE project website. Interacting pairs of genomic regions from this files corresponds to two nodes linked by an edge in our network. The data corresponding to two different ChIA-PET replicates were downloaded and only interactions supported by both replicates were retained for further analysis.

#### Filtering of PET interactions

As we were interested in *cis* long-range interactions we filtered out inter-chromosomal PET interactions before further analysis. Also we excluded the so-called self-ligation PETs from further analysis ([20]). They represent an artifact of ChIA-PET experiments, and originate from self-circularization ligation of the same chromatin fragment resulting in ChIA-PET sequences with both tags mapped within a short genomic distance of each other. In order to distinguish between self-ligation PETs and inter-ligations PETs, which actually correspond to two distinct interacting chromosomal regions, we performed a similar analysis to Li et al. ([20]). We computed the genomic distances between PETs and plotted their frequency in each genomic bin on a log-log scale. The intersection of two fitted lines at 1691 nt was taken as distance cutoff to distinguish self-ligation from inter-ligation PETs, which seem to follow two distinct power-law distributions (Fig. 2). According to this cutoff, self-ligation interactions, with distances below this cutoff, were discarded from further analysis.

**Figure 2:**
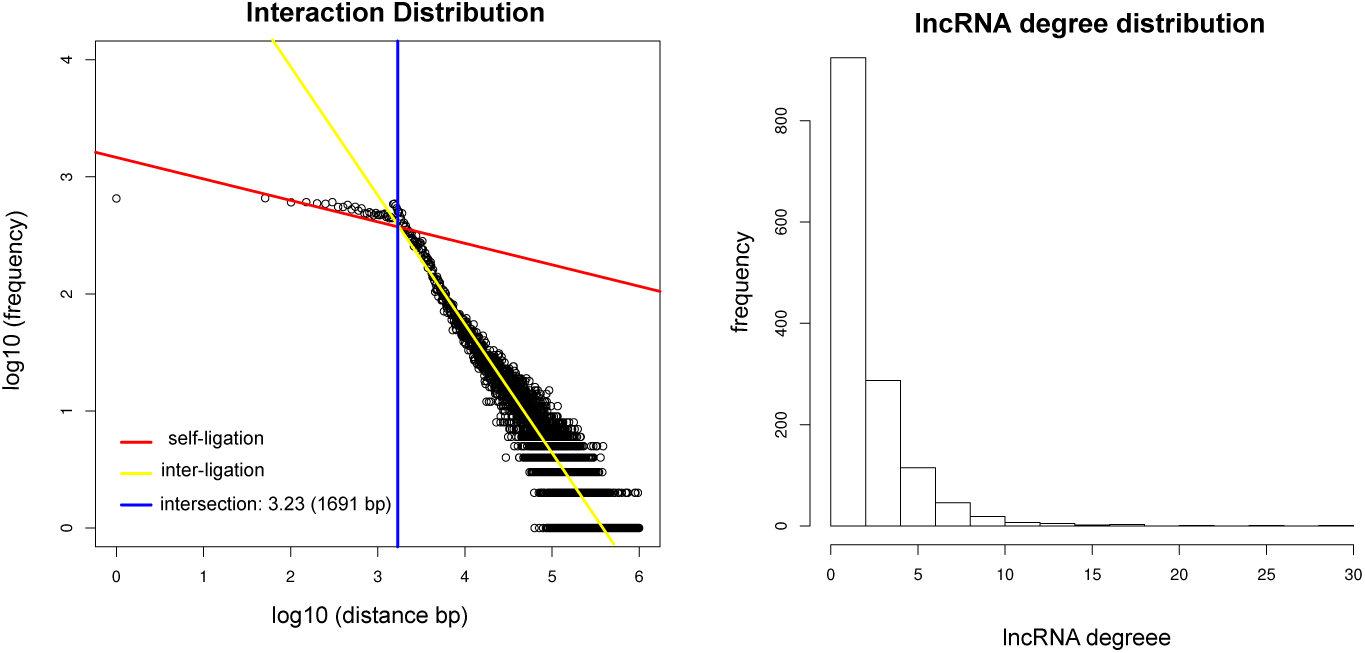
Filtering of interacting regions. Left panel: Fitted mixture model to classify PETS in self-ligation and inter-ligation. Rigth panel: Degree centrality distribution for lncRNAs

#### Expression analysis of lncRNAs and genes

Only lncRNAs and genes detected at a certain expression level in the K562 RNA-seq data were used in the subsequent network analysis. For both genes and lncRNAs, alignment files in K562 were obtained from the Cshl Long RNA-Sequencing track on the ENCODE website. Genomic annotation of lncRNAs and genes was taken from Gencode v24. Gene and lncRNA coordinates were lifted over the hg19 human genome assembly as all other annotations were on hg19. Read counts in genes and lncRNAs were obtained by means of htseq-count for two different replicates and converted to RPKM values. To quantify the final expression level of a gene or a lncRNA, RPKM values between the two replicates were averaged. Only detected lncRNAs (RPKM *>*0) and genes with an RPKM value higher than 0.071 were included in the network construction. This latter threshold was determined by looking at the bimodal distribution of the log RPKM expression values and corresponds to the local minimum separating the two modes.

#### Network construction and annotation

PETs representing interacting genomic regions were annotated as ‘gene’, and assigned their corresponding official gene symbol, if they overlapped the genomic coordinates of annotated genes from Gencode. PETs were annotated as ‘lncRNA’ if they overlapped the genomic coordinates of annotated lncRNAs from Gencode. Given that the resolution of the ChIA-PET data is in the order of few kilokases, it could occur that interacting PETs might cover wide genomic regions with more than one annotated gene/lncRNA. In addition, ChIA-PET data are not strand-specific, therefore they might overlap with two or more genes/lncRNAs located on different strands. PETs corresponding to more than one gene/lncRNA location, either on the same or the opposite strand, were annotated with both gene and lncRNA names. Chromatin states in K562 from the the chromHMM genome segmentation [22] downloaded from the ENCODE website were also used to annotate interacting PETs in the network. ChromHMM uses a hidden Markov model to annotate genomic regions according to combinations of chromatin modifications and Transcription Factor Binding Sites (TFBSs). This results in a segmentation of the genome in ‘enhancer’, ‘weak enhancer’, ‘TSS’, ‘promoter flanking’, ‘CTCF’, ‘transcribed’ and ‘repressed’. The assignment ‘repressed’ was ignored because in a network containing interactions mediated by PolII, repressed regions hold no information. PETs were annotated with chromHMM categories if they overlapped the genome segmentation annotation (Fig. 1b). It could occur that the same PET overlapped with many different features. In this case annotations were merged. For example a PET overlapping both an annotated lncRNA and an enhancer region was defined as ‘lncRNA enhancer’. All interacting PETs were assigned a unique identifier. If PETs did not overlap with any annotated gene, lncRNA or chromHMM state they were labeled as *unknown*. Annotated PETs were represented as nodes in the network and an interaction between PETs as an edge. A global (0,1)-adjacency matrix was build to describe the overall graph, called from now on *chromatin graph*. In addition, an adjacency matrix or individual graph was created separately for each chromosome. The number of rows and columns of the adjacency matrix represents the number of genomic regions involved in at least one ChIAPET interaction. A 0-entry in the matrix cell corresponds to no interactions between any two PETs overlapping with these regions, while a 1-entry corresponds to a ChIA-PET interaction. A schematic view of the steps described above is given in Fig. 1b. Next to their name, type and expression, genes and lncRNAs were also annotated with associated diseases and information about whether they are part of a positionally conserved gene - lncRNA pair. Disease annotation data was taken from the database lncRNADisease (as of June 2015) [23] where we used both experimentally validated associations between lncRNAs and diseases, as well as predicted associations. LncRNAs that were part of positionally conserved pairs of genes and lncRNAs were obtained from [24].

### Network analysis of the chromatin graph

#### Centrality measures

For graph analysis we use standard graph concepts of interest for biological network analysis, see, e.g., [25] and [17]. To identify nodes of potential functional importance, we first look for nodes with a high degree, i.e., with a high number of incident edges. In biological networks that have a scale-free structure, such nodes are called hubs and they often correspond to elements of biological interest, since they interact with a large number of other nodes. For each node *v* in a graph *G* = (*V, E*) we calculate the number *d*(*v*) of edges incident to *v* and call it its degree or degree centrality. Another node measure of interest is a betweenness centrality, which captures the importance of a node as an efficient connector to other nodes in the network. A node may have a rather low degree, but still carry great importance for network functionality if connecting otherwise rather isolated clusters or, modules of the graph. For *v* ∈ *V*, betweenness centrality is defined as *b*(*v*) = ∑_*s≠v≠t*_ (*σ*_*st*_(*v*)*/σ*_*st*_), where *σ*_*st*_ is the number of shortest paths from node *s* to node *t* and *σ*_*st*_(*v*) is the number of such paths that pass through *v*.

#### MSM clustering for module detection

The graphs we are interested in are often not connected, i.e., there exist nodes in the network that cannot be connected via a path. We then focus on the connected components of the graph. Formally, a connected component *C* = (*V*_*C*_, *E*_*C*_) is a maximal subgraph of *G* such that there exists a path between *v* and *w* for all vertices *v, w ∈ V*_*C*_. Given a connected graph or connected component, we are interested to identify substructures, in particular, subgraphs that have a high connectivity but are only sparsely connected to the rest of the network. These graph structures are called modules and in biological networks they often represent functional units.

In this paper, we apply the MSM clustering method developed in [18, 19], which is based on finding Markov State Models (MSM) of a time-continuous random walk process. MSM clustering is a dynamics-based method, which uses properties of the random walk process to discover the network structure. More precisely, it identifies modules as regions of the network where the process is metastable, i.e. trapped for a longer period of time. To this end, the number of network modules can be induced from the number of dominant eigenvalues of the generator matrix that governs the dynamics of the random walk process. Unlike most of the common approaches that can identify only complete partitions of the network, MSM finds fuzzy partitioning into modules. Namely, with MSM we can identify modules *M*_1_, *…, M*_*m*_ and a transition region 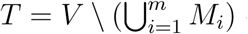 consisting of nodes which are not uniquely assigned to exactly one of the modules, but can belong to several modules or to none. This makes the approach particularly suited to biological applications where some molecular components may have multiple functions or are not necessarily integrated in functional units. For every node *x* we can calculate a value *q*_*i*_(*x*) as the random walk based probability of affiliation of a node *x* to a module *M*_*i*_. We then use a free parameter *θ* to refine the partitioning, i.e. we assign a node *x* to a module *M*_*i*_ if *q*_*i*_(*x*) *≥ θ*. If *θ* = 1 we obtain the core modules, meaning the subgraphs exhibiting the strongest cohesiveness w.r.t. the module characteristics. By decreasing *θ* we expand modules until we reach a full partitioning of a graph by associating each vertex from the transition region with exactly one module it most likely belongs to. Fuzzy affiliation functions *q*_*i*_, *i* = 1, *…, m* can be obtained by solving sparse, symmetric and positive definite linear systems ([26, 18]).

Another free parameter is a resolution parameter *α*, indicating how densely connected the modules we are interested in finding should be. That is, for high values of *α,* the method finds dominant, highly intraconnected modules and by decreasing *α* it finds also less pronounced modules. This is connected to the timescale at which the random walk leaves the transition region. It can be originally set according to the gap in the dominant spectrum of the generator of the random walk and then varied to observe the effect on the modules. In our application, it usually ranges from 100 to 2000.

#### Optimization criteria

The parameters *θ* and *α* allow for an adaptation of the clustering to the specific application. Since we are looking for regulatory units involving lncRNAs, we chose to compare co-expression levels of intraversus inter-modular gene-lncRNA pairs in order to find the best clustering parametrization and to assess the quality of the clustering in general. We argue that elements within the same module should have more correlated expression profiles, indicating potential mutual regulation, whereas intermodular node pairs are more independently regulated. In detail, we performed the MSM clustering for connected components from all chromosome graphs for a range of *α* and *θ* combinations. We chose the best combination by optimizing an empirical objective function (Equ.1) defined by the ratio of the median intra-module mutual information (MI) and the inter-module MI for all gene-lncRNA pairs in the connected component.

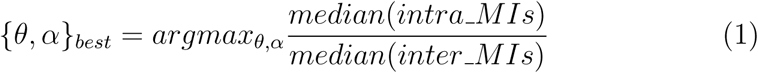

MI values for gene-lncRNA pairs were computed based on RKPM expression values (taken from ENCODE RNA-seq, see above) across 34 different tissues. First the optimal binning of each vector of continuous values is found and then the marginal entropies of the two distributions minus their joint entropy are computed, as described in [27]. The reported ratio in Equ.1 for a connected component serves also as indicator for the quality of the clustering, where a high score implies a better partitioning with respect to MI and a ratio of at least one is expected for biologically meaningful clusterings.

### Comparison with other clustering methods

We compared our MSM clustering approach to other state-of-the-art clustering methods with respect to the mutual information ratio capturing co-expression within the calculated modules. We used the following methods and their implementation from the R *igraph* package [28]:

- *cluster fast greedy* function (FG), which finds dense subgraphs by directly optimizing a modularity score *Q*. Given a set of modules, *Q* is computed as the ratio between the fraction of within-community edges versus the expected fraction of connections for the randomized network [29].
- clustering via edge betweenness (EB), *cluster edge betweenness* function, which is based on iteratively removing edges with highest edge betweenness from the graph [30], in order to hierarchically split the graph into modules.
- leading eigenvalue clustering algorithm (EV), *cluster leading eigen* function, which implements the popular graph clustering method from Newman [31]. Instead of directly optimizing the modularity score *Q*, this method finds network modules by calculating the leading non-negative eigenvector of the so called modularity matrix.
- Walktrap algorithm which is a random walk based clustering (RRW), *cluster walktrap* function. Similarly to our MSM algorithm this approach finds modules in a graph by exploiting metastability of the random walk [32], but uses only a time-discrete version of the process.

Unlike our method, all aforementioned clustering approaches do not support fuzzy clustering. So we compare these methods to our MSM procedure over a large set of method-specific parameters for the largest connected component of our chromatin graph on chromosome 1. This comparison is not straight-forward since the modularity score *Q* which most of these methods use is hard to compare between fuzzy and non-fuzzy clustering and might not be very meaningful in our context. In addition, we optimize the parameters of our MSM determining size and number of modules through the MI ratio defined above, therefore a similar evaluation should be done for the above mentioned algorithms.

To address these issues, we evaluated a range of different modules for each of the considered methods from the *igraph* package. First, we assessed the modules of each method returned by their own underlying optimization algorithm. As additional information to this clustering, most of the considered algorithms return a hierarchical overview of the best clusterings for a range of different module numbers - comparable with the variation of the parameters of MSM - which can be accessed via the so-called *cut at* function. This allows us then to assess the results for clusterings corresponding to slightly larger resp./ smaller module number (+/- 1) than the number of modules of the optimal clustering. If the change in number of modules resulted in a change of MI ratio, we continued increasing or decreasing the number of modules incremently and evaluated the resulting modules (figure 4 gives an impression of the magnitude of change). As a last step, we considered the results of the different methods obtained for the optimal module number generated by our MSM clustering to gain more information about the quality of that number. An exception to this procedure is the EV algorithm that does not offer a simple way to change the number of modules. Rather, we can only influence this number indirectly using the ‘steps’ parameter, which allows only for small variations in the number of modules. The EB algorithm, for example, finds 19 modules in chromosome 1. We then produced clusterings with 17, 18, 20 and 21 modules as well as with 11 modules (the same as for the MSM method).

**Figure 3:**
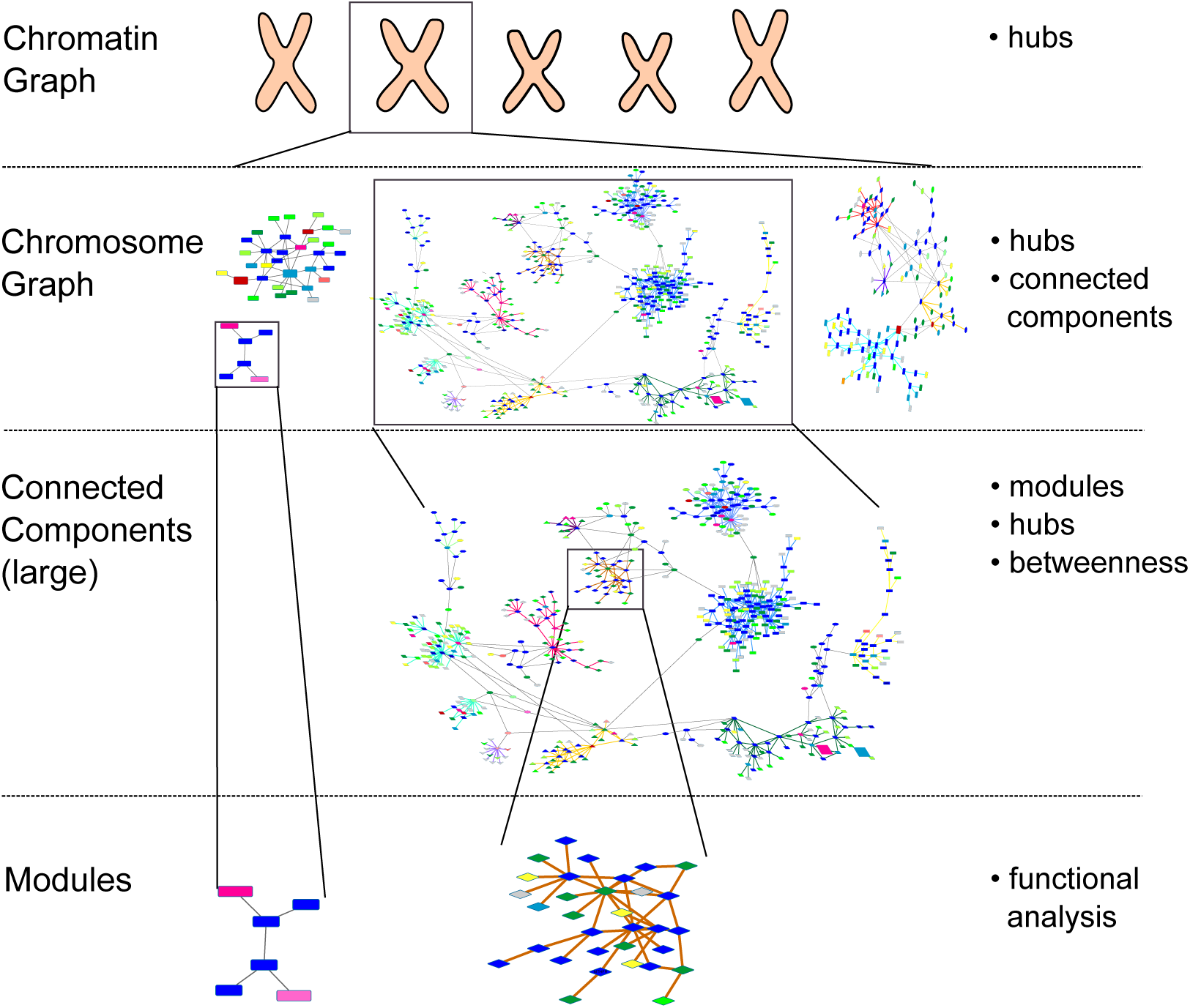
Overview of the hierarchical graph analysis. On the right are listed the different analysis steps executed for each level. Both small connected components as well as density-based modules of large connected components are found on the lowest level.

**Figure 4:**
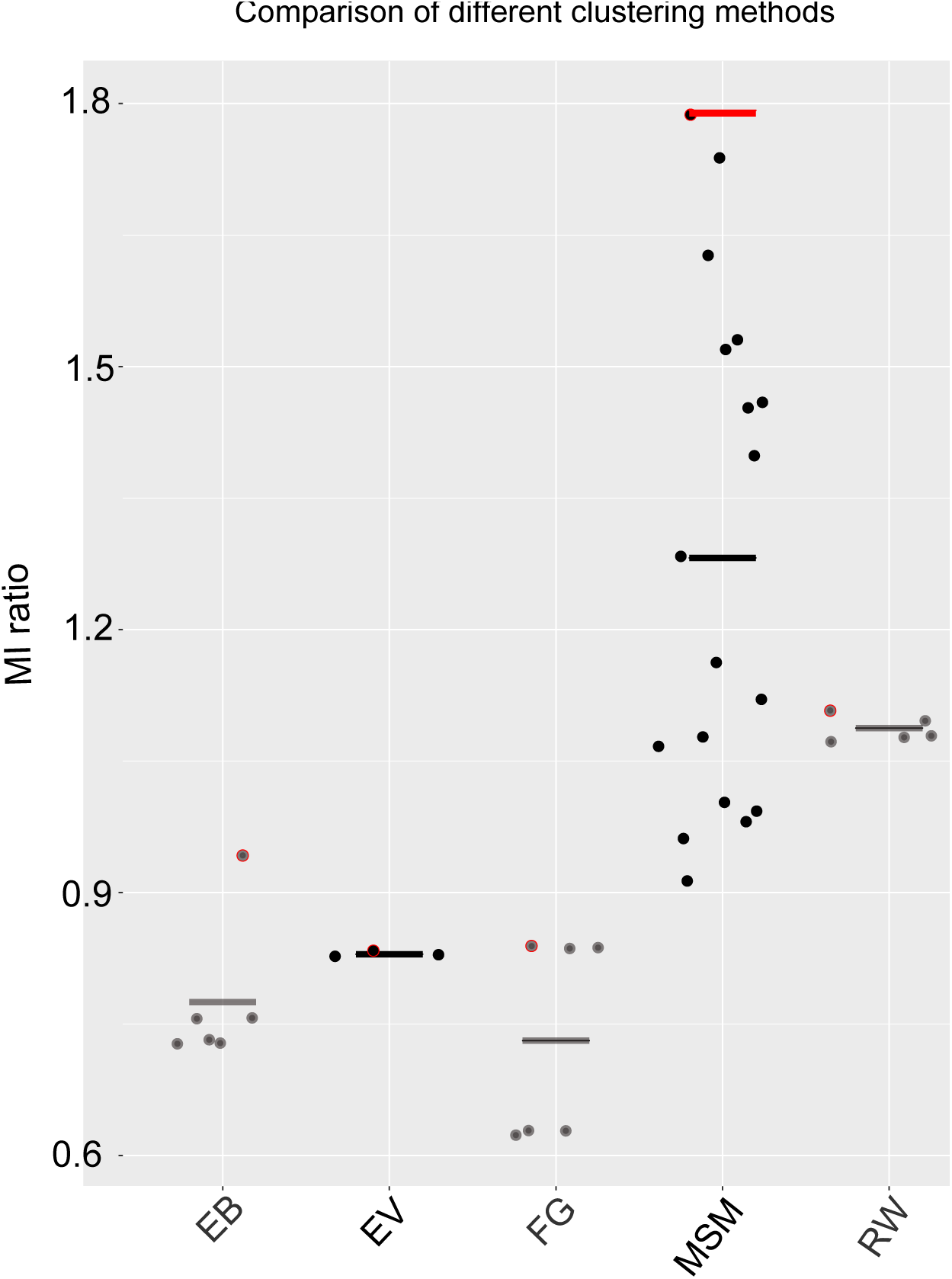
Comparison of different graph clustering methods. Our MSM clustering approach is compared to other methods from the *igraph* package (EB - clustering via edge betweenness; EV-eigenvalue clustering; FG-fast and greedy clustering; RW-random walk clustering). All methods are run with different ranges of parameters and/or number of modules, and the mutual information (MI) ratio is computed for every scenario as described in Material and Methods. For each method the distribution of the resulting MI ratio is shown, together with the median value (horizontal line). For each clustering method the result obtained with the MSM’s optimal number of modules is circled in red. The red line indicates the best partition for our MSM clustering, i.e. values of *α* and *θ* yielding the highest MI ratio.

## Results

In this section we present the results from the network construction, the features of the chromatin graph, as well as connected components for each chromosome. We focus on the analysis of different centrality measures for lncRNA nodes and their relation to other genes or regulatory elements. In the last paragraph we discuss some important lncRNA-mediated functional modules identified by means of the MSM clustering method described above.

### Hierarchical graph analysis of the ChIA-PET interaction network

The genomic distance cutoff for distinguishing ChIA-PET interactions into short range (i.e. self-ligation) and long-range (i.e. inter-ligation) interactions was derived from the log-log plot displaying the frequency of interactions at different genomic distances (Figure 2, Left panel). The plot clearly shows two linear ‘regimes’, corresponding to a mixture distribution of PETs where two different linear functions can be fitted. The intersection of the two fitted lines in the log-log plot was chosen as cutoff to differentiate self-ligation from inter-ligation. Self-ligation PETs were excluded from the network analysis as, in most of the cases, they do not correspond to chromatin interactions between different genomic segments.

To cope with the size and heterogeneous nature of the chromatin graph we developed an hierarchical analysis approach that allows to add step-wise resolution to subgraphs of interest guided by the results of the previous step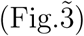. We start by analyzing the global chromatin graph to identify global hubs, computing the degree centrality for each lncRNA in the chromatin graph. Additionally the *to gene degree*, meaning the number of neighboring genes for each lncRNA, was extracted to distinguish more important interactions. With these two measures an enrichment analysis was conducted (described in detail in the next two paragraphs) to understand the properties of hub lncRNAs with respect to disease, positional conservation and enhancer annotation. An overview of the general properties of the chromatin graph is given in Table 1. For each chromosome we computed: the number of connected components, the minimum, average and maximum size of the components, the total number of annotated lncRNAs, the total number of lncRNAs involved in ChIA-PET interactions, and report the lncRNA with highest degree. As it can be noted from Table 1, the chromatin network is very sparse, with many components representing singleton nodes or containing very few nodes. When looking at components of bigger size, we notice that only few lncRNAs have a degree centrality higher than 10, while the ma-jority of lncRNAs exhibits a degree between one and three. This matches the general observation that in biological networks degrees are often distributed according to a power law, i.e., there exist few hubs and many much less densely connected nodes [17]. We ranked lncRNAs according to their degree, in order to identify global hubs of the chromatin graph. Table 2 reports the top 20 highly-connected lncRNAs from the network, which have RPKM *>* 0.

**Table 1:**
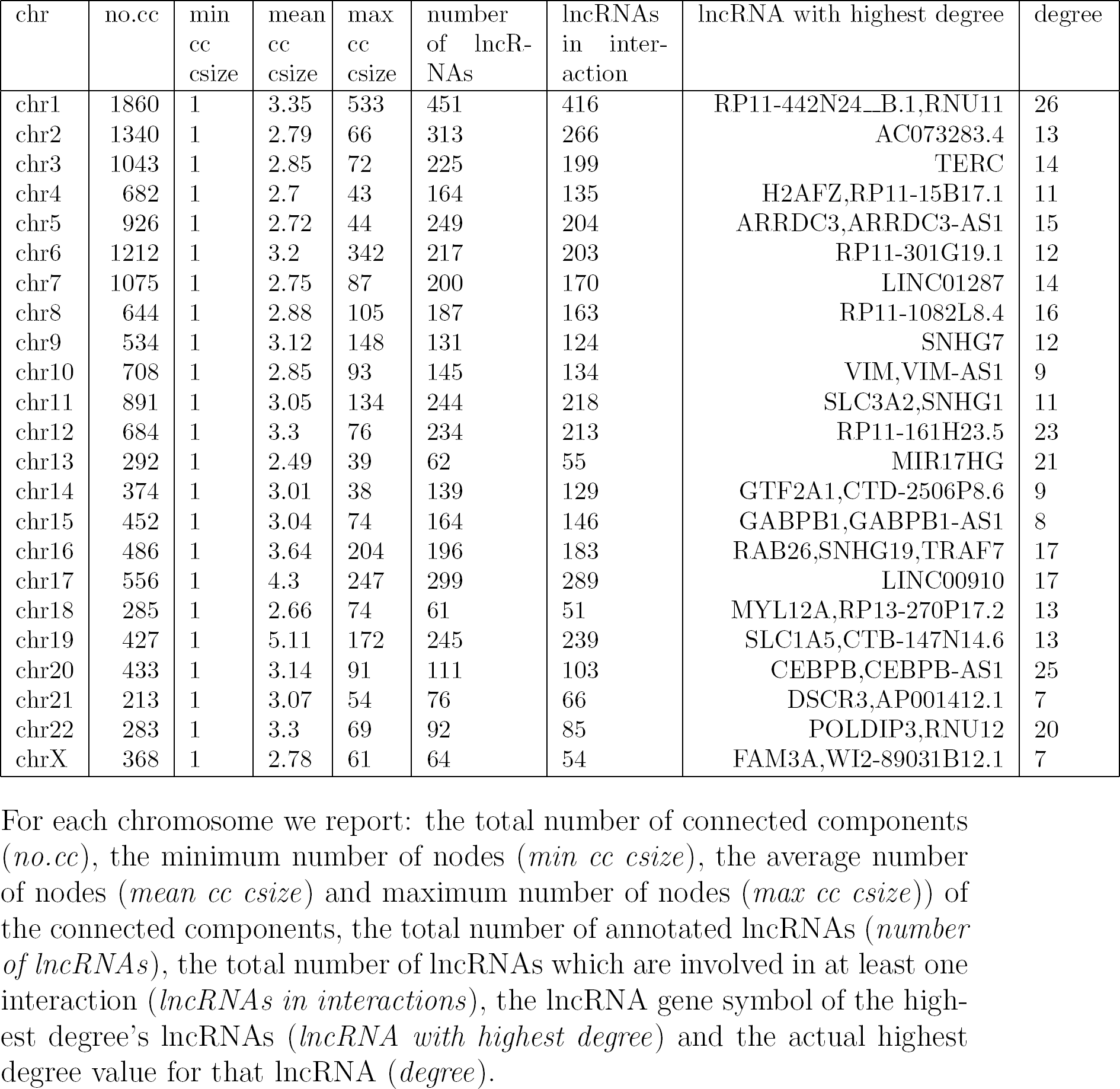
Properties of the chromatin graph

| chr | no.cc | min<br>cc<br>csize | mean<br>cc<br>csize | max<br>cc<br>csize | number<br>of lncR-NAs | lncRNAs<br>in inter-<br>action | lncRNA with highest degree | degree |
| --- | --- | --- | --- | --- | --- | --- | --- | --- |
| chr1 | 1860 | 1 | 3.35 | 533 | 451 | 416 | RP11-442N24_B.1,RNU11 | 26 |
| chr2 | 1340 | 1 | 2.79 | 66 | 313 | 266 | AC073283.4 | 13 |
| chr3 | 1043 | 1 | 2.85 | 72 | 225 | 199 | TERC | 14 |
| chr4 | 682 | 1 | 2.7 | 43 | 164 | 135 | H2AFZ,RP11-15B17.1 | 11 |
| chr5 | 926 | 1 | 2.72 | 44 | 249 | 204 | ARRDC3,ARRDC3-AS1 | 15 |
| chr6 | 1212 | 1 | 3.2 | 342 | 217 | 203 | RP11-301G19.1 | 12 |
| chr7 | 1075 | 1 | 2.75 | 87 | 200 | 170 | LINC01287 | 14 |
| chr8 | 644 | 1 | 2.88 | 105 | 187 | 163 | RP11-1082L8.4 | 16 |
| chr9 | 534 | 1 | 3.12 | 148 | 131 | 124 | SNHG7 | 12 |
| chr10 | 708 | 1 | 2.85 | 93 | 145 | 134 | VIM,VIM-AS1 | 9 |
| chr11 | 891 | 1 | 3.05 | 134 | 244 | 218 | SLC3A2,SNHG1 | 11 |
| chr12 | 684 | 1 | 3.3 | 76 | 234 | 213 | RP11-161H23.5 | 23 |
| chr13 | 292 | 1 | 2.49 | 39 | 62 | 55 | MIR17HG | 21 |
| chr14 | 374 | 1 | 3.01 | 38 | 139 | 129 | GTF2A1,CTD-2506P8.6 | 9 |
| chr15 | 452 | 1 | 3.04 | 74 | 164 | 146 | GABPB1,GABPB1-AS1 | 8 |
| chr16 | 486 | 1 | 3.64 | 204 | 196 | 183 | RAB26,SNHG19,TRAF7 | 17 |
| chr17 | 556 | 1 | 4.3 | 247 | 299 | 289 | LINC00910 | 17 |
| chr18 | 285 | 1 | 2.66 | 74 | 61 | 51 | MYL12A,RP13-270P17.2 | 13 |
| chr19 | 427 | 1 | 5.11 | 172 | 245 | 239 | SLC1A5,CTB-147N14.6 | 13 |
| chr20 | 433 | 1 | 3.14 | 91 | 111 | 103 | CEBPB,CEBPB-AS1 | 25 |
| chr21 | 213 | 1 | 3.07 | 54 | 76 | 66 | DSCR3,AP001412.1 | 7 |
| chr22 | 283 | 1 | 3.3 | 69 | 92 | 85 | POLDIP3,RNU12 | 20 |
| chrX | 368 | 1 | 2.78 | 61 | 64 | 54 | FAM3A,WI2-89031B12.1 | 7 |

For each chromosome we report: the total number of connected components (*no.cc*), the minimum number of nodes (*min cc csize*), the average number of nodes (*mean cc csize*) and maximum number of nodes (*max cc csize*) of the connected components, the total number of annotated lncRNAs (*number of lncRNAs*), the total number of lncRNAs which are involved in at least one interaction (*lncRNAs in interactions*), the lncRNA gene symbol of the highest degree's lncRNAs (*lncRNA with highest degree*) and the actual highest degree value for that lncRNA (*degree*).

**Table 2:**
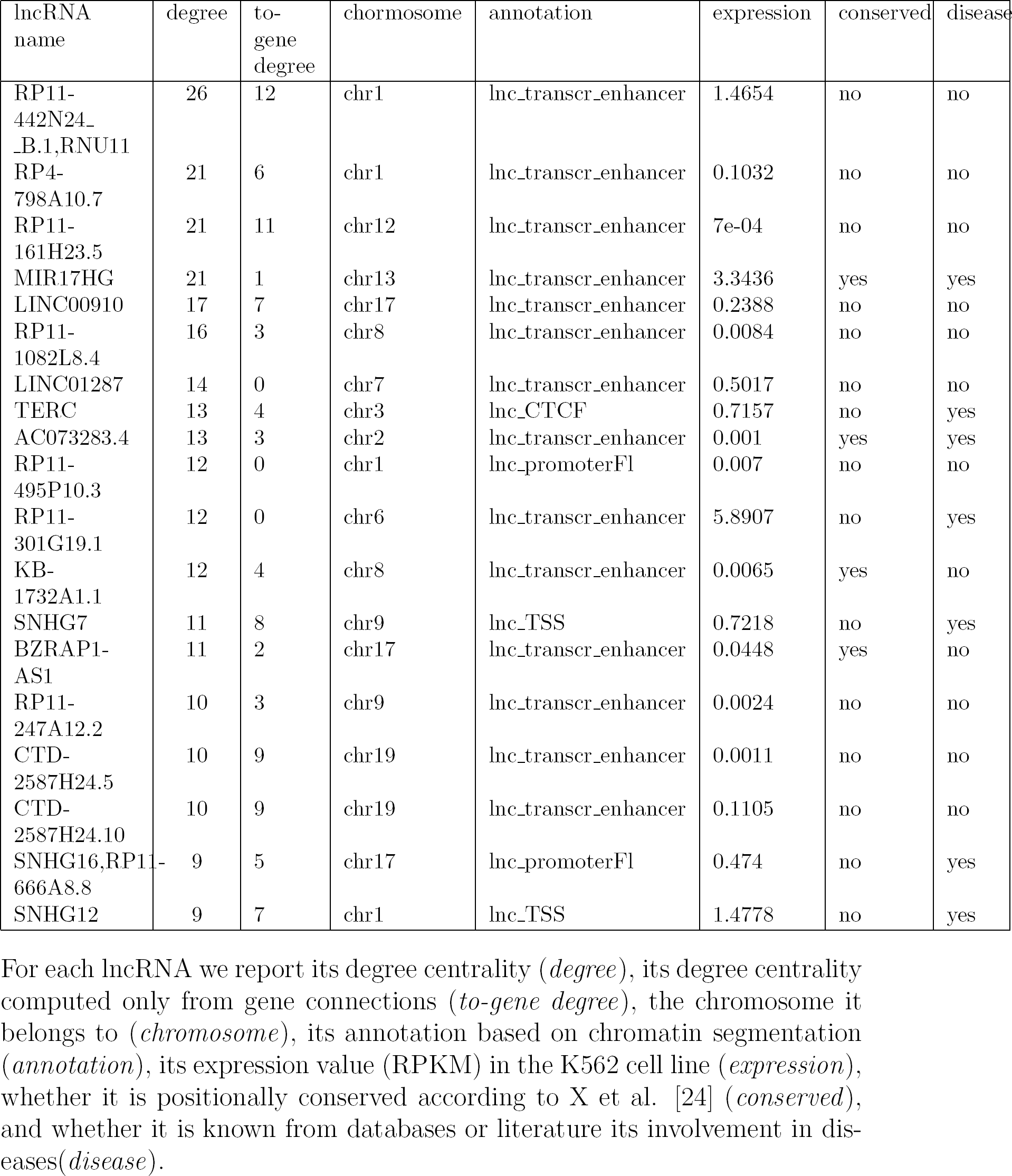
Top 20 lncRNAs with highest degree from the chromatin graph.

Since the chromatin graph decomposes in a natural way into the graphs representing the single chromosomes, we consider the lncRNA nodes within the respective isolated chromosome graphs next and compute the lncRNA degree chromosome-wise. Even nodes that are not among those of highest degree in the chromatin graph may be distinguished with respect to their chromosome graph. Second, we focus on the connected components containing lncRNAs of each chromosome graph to obtain the next resolution level. Small components are then amenable to a full analysis of different aspects of interest, including node centralities, but also subgraph composition and topology w.r.t. the node annotations. For large connected components we still need indicators that guide our search for important lncRNA modules. As on the chromosome level, we repeat the degree analysis to get results relative to the topology of the connected component. In Tables S1, S2 and S3 (Additional file 1) we report this analysis for the biggest connected components of chromosome 1, 17 and 11, respectively. In addition, since when we look at connected components we deal with a connected graph, we evaluate the betweenness centrality of each lncRNA node. Such nodes can have great importance for mediating between the majority of the network nodes, different clusters or modules in the graph. To uncover a possible structuring of the connected component and identify relevant functional units, we conduct a module search using the Markov State Model (MSM) clustering method described above. As mentioned, depending on the parametrization of the method, the algorithm searches for highly intra-connected modules.

### LncRNAs with high degree-centrality are associated to diseases

By manually inspecting the functional annotation of the top 20 expressed lncRNAs with highest degree, we find several lncRNAs known from previous studies to be cancer-associated. For example, RNAs from the SNHG family, such as SNHG16, SNHG7 and SNHG12, are lncRNAs which are host genes of small nucleolar RNAs, known to be important in cell proliferation and invasion in different cancer types, such as osteosarcoma, lung cancer and others [33]. The lncRNAs RP11-301G19.1 has been found to be over-expressed in leukemia [34]; TERC, a lncRNA ivolved in telomerase activity has been strongly associated to leukemic cells [35], and the intergenic lncRNA MIR17HG, host transcript of the MiR-17-92a-1 cluster is known to be involved in cell survival and cancer proliferation [36]. However, disease annotation is sparse and limited for lncRNAs compared to protein-coding genes. The fraction of intergenic lncRNAs from the ChIA-PET network, that could be annotated with a disease in our analysis (see Matherial and Methods for detail) was only 6% (188 out of 3172), therefore it is hard to systematically access whether high-degree lncRNAs are significantly associated to disease, and specifically to cancer. Comparing the degree distribution of intergenic lncRNAs (lincRNAs) annotated with a disease versus lincRNAs not linked to a disease we observe a global trend, although not statistically significant, of high-degree lincRNAs to be disease-associated (p-value 0.207, Wilcoxon rank sum test). When we perform the same analysis by including in our lncRNA dataset not only intergenic lncRNAs, but also lncRNAs overlapping protein-coding genes on the opposite strand (i.e. antisense lncRNAs), we can assign a disease annotation up to 9% of the lncRNAs in our network, and obtain a slightly significant association between degree centrality and disease annotation (p-value 0.086, Wilcoxon rank sum test).

### Positional conservation and enhancer functions of high-degree lncRNAs

Further analysis of lncRNAs not overlapping with gene regions showed that lncRNAs annotated as “transcribed enhancers”, according to chromHMM predictions, have a significantly higher degree centrality compared to those lncRNAs annotated as “non-transcribed enhancers” (p-value=1.9 *\** 10^−19^, Wilcoxon rank sum test). This interesting observation points to the important role of the act of transcription for loop-mediated transcriptional regulation. In addition, when we compare lncRNAs, which are also annotated as enhancers from chromHMM (independently from being transcribed or not), to lncRNAs which do not overlap enhancer elements, we find that enhancer-like lncRNAs have significantly higher degree compared non-enhancer lncRNAs (p-value=3.2 *\** 10^−65^, Wilcoxon rank sum test). This suggests that enhancer-like lncRNAs are hubs in the PolII-mediated ChIA-PET gene regulatory network and exert their regulatory role by connecting several, putatively functional genomic regions to gene loci in an extensive and combinatorial fashion.

Previous studies have hypothesized that, in the absence of extensive sequence conservation, “positional” conservation of lncRNAs between human and mouse, meaning that their genomic position is preserved relative to orthologous coding genes, are potentially functional [24]. Positionally-conserved lncRNAs have also been associated with developmental or cancer genes, and shown to be in chromatin loops, which contact enhancer-regulatory sequence. In our network, we observe that lncRNAs which are also annotated as positionally conserved, have a significantly higher degree than not positionally conserved ones (p-value=1.9*\**10^−19^, Wilcoxon rank sum test), indicating their potential role as functional hubs in the PolII chromatin network.

### MSM module detection and comparison with existing strategies

We apply our MSM clustering method to detect potential regulatory modules, i.e. groups of highly interconnected genes, lncRNAs and different chromatin states in the K562 cell line for each chromosome’s biggest component. As explained in Material and Methods, we do that by selecting the *α* and *θ* parameters of the clustering algorithm which maximize the ratio of the intra-cluster mutual information (MI) versus the inter-cluster MI. Pairwise MI values represent the correlation in expression between lncRNAs and their associated targets across ENCODE cell lines. Therefore our empirical objective function reflects the expectation that lncRNA-gene modules showing a high density of chromatin-connected regions have correlated expression profiles. The best values for *α* and *θ* for each inspected connected component are reported in the table of Additional file 2, which also lists the optimal number of clusters detected for each component, the number of nodes in each cluster which are associated to either a protein coding gene or a lncRNA, and therefore represent putative regulatory modules. By analyzing each chromosome’s biggest component (Additional file 2) we observe that generally clusterings with *θ* = 0.7 and small *α* (around 100-500), allowing more sparsely connected and relaxed modules, provide the highest MI ratio. Other parameters are mostly found when networks get too small and do not provide enough nodes with expression levels. We compared the results of our MSM approach to several state-of-the-art methods, as described in the Method’s section. Our method clearly outperforms all of these clustering methods when evaluating the mutual information and Figure 4 shows that even the mean clustering result from MSM is higher than the best clustering obtained from other methods. It is noteworthy that the best clusterings of other methods are the ones with the number of modules reported by MSM. Among these methods, the Walktrap algorithm delivers the best results, which indicates that random-walk-based methods are better suited for the complex structure of the networks we are dealing with.

### Biologically-relevant detected regulatory modules

Prior to cluster analysis, we first inspected small connected components of the network (containing less than 50 nodes). Fig. S1a) (Additional file 1) shows the module containing the lncRNA CYP4A22-AS1, also known as ncRNA-a3, which has been shown to act as enhancer for its flanking stem cell leukemia-associated gene TAL1 [10]. Looking at the network connections of this component we recapitulate the direct interaction between ncRNA-a3 and TAL1, putatively mediated by STIL, a regulator of mitotic spindle involved in T cell leukemias. The active enhancer-like lncRNA linc00853, also known as ncRNA-a4 is also part of the ncRNA-a3 network (Fig.S1a)) and directly regulates its flanking gene CMPK1, as already previously verified experimentally [10]. Interestingly, this suggests a synergistic action of these two lncRNAs in coordinating the transcriptional activity of a group of four genes in this module connected to the two lncRNAs at chromatin level.

We also looked at the well characterized lncRNA Xist (Additional file 1: Fig. S1b)), known to be one of the main drivers of transcriptional gene silencing during X-chromosome inactivation, but whose mechanisms of action are still not fully understood. From our analysis it is evident, given the lack of ChIA-PET interactions, that Xist does not associate to PolII to regulate its target genes in an enhancer-like fashion, in agreement with its suggested silencing function. On the other hand, we could recover direct PolII-mediated interactions between XIST and lncRNA FTX and JPX, which are known regulators of XIST transcription, as well as a possible interaction with TSIX Fig.S1b) which is located anti-sense to XIST [8].

In Fig. 5 we also show the results of our clustering analysis on three selected biggest components from chromosome 1, 11 and 17, respectively, where modules are defined on the basis of the best combination of *α* and *θ* parameters, as described in Materials and Methods.

**Figure 5:**
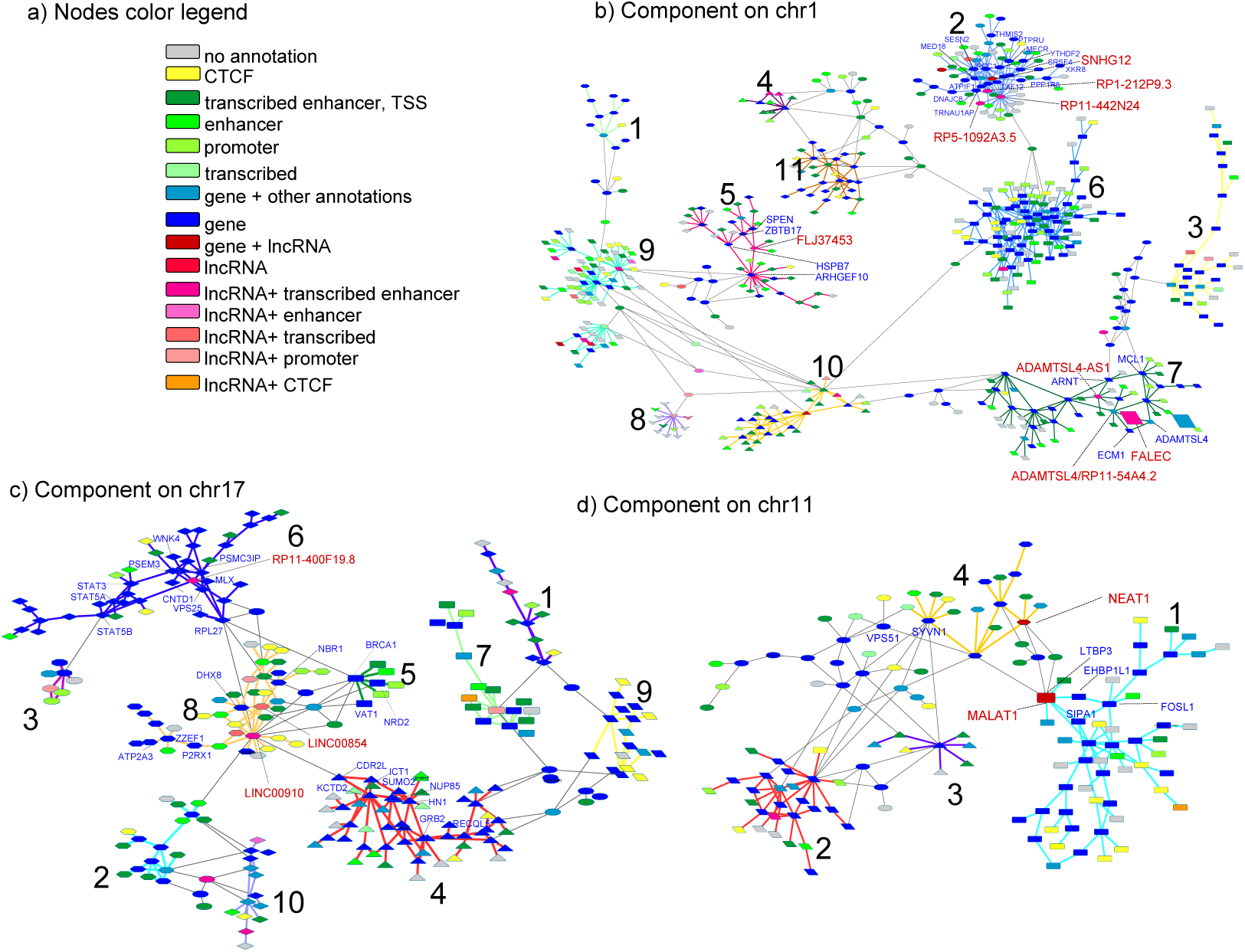
Module detection. a) Node color legend, b) chromosome 1 biggest component, c) chromosome 17 biggest component and d) chromosome 11 biggest component. Nodes are colored according to their overlaid genomic annotation and/or chromatin states (a). Relevant lncRNAs and connected genes are highlighted as text. Different modules from the clustering analysis are represented via different nodes’ shapes and edges’ colors. ‘Fuzzy’ nodes, i.e. nodes which could not be confidently assigned to any module, have connections colored in gray departing from them.

### Module structure and putative lncRNA functions in chromosome 1’s biggest component

We detect 11 modules for the biggest component on chromosome 1 obtained with setting *α* = 1000 and *θ* = 0.9 (Fig. 5c). Nodes belonging to different clusters are depicted in different shapes and different clusters are also recognizable by different colors of their intra-cluster edges. Highlighted in Fig 5c is the regulatory module encompassing 57 interacting chromatin regions (nodes) and containing the well-known eRNA FALEC (cluster 7), which has been shown to have enhancer-like functions and significantly influence the expression of its flanking gene ECM1 [10], also contained in the same module. By inspecting the interactions in this module, we learn that the interaction between FALEC and the ECM1 locus is mediated by the ADAMTSL gene and the lncRNA RP11-54A4.2. Interestingly, the Myeloid Cell Leukemia apoptosis regulator MCL1 and the Hypoxia-Inducible factor 1-Beta ARNT are also in the same module, indirectly linked to FALEC via other lncRNAs, protein-coding genes and several transcribed enhancer elements (colored in green in Fig. 5c). These results suggest an involvement of FALEC in the regulation of these cancer genes in the leukemia cell line, although additional validation experiments would be necessary to show an effect of FALEC on the distant genes ARNT and MCL1.

Of note is a modular unit on chromosome 1 containing the lncRNA of unknown function FLJ37453 (cluster 5), which connects genes such as the transcriptional repressors ZBTB1 and SPEN and the tumor suppressor HSPB7. While one could hypothesize that FLJ37453 might regulate the activity of HSPB7, the presence of SPEN in this cluster is interesting. The hormone inducible repressor SPEN has been shown to associate with Xist lncRNA to silence gene transcription through the recruitment of histone deacetylases or the sequestration of transcriptional activators [37]. Here, the physical association of FLJ37453 lncRNA with SPEN on chromatin could indicate a co-regulation mechanisms necessary to orchestrate a coordinated transcriptional response and a putative role of FLJ37453 in SPEN-mediated gene silencing.

We also examined cluster 2, the module on chr1 containing, among others, lncRNA RP11-442N24, the highest degree node of this connected component. Interestingly, GO analysis revealed that this cluster contains genes enriched in GO terms “RNA PolII transcriptional pre-initiation complex”, “histone acetylation”, “SLIK (SAGA-like) complex” and “TF IID complex” (corrected p-value*<*= 0.1). This suggest a role of RP11-442N24 in general transcriptional regulation and chromatin remodeling by recruiting transcription factors of the transcriptional pre-initiation complex, mainly involved in histone acetylation.

### Module structure and putative lncRNA functions in chromosome 11’s biggest component

Clustering of the biggest component on chromsome 11 (Fig. 5d) results in an optimal partition of four dominant modules, obtained with setting *α* = 500 and *θ* = 0.7 (thus identifying more relaxed modules). Particularly interesting are the modules marked by yellow and blue connections, namely cluster 1 and 4, linked by the oncogene lncRNA MALAT1, known to act as transcriptional regulator for numerous genes involved in cancer metastasis and cell migration [6]. MALAT1 has a degree centrality value of 8 in its component, but exhibits a very high betweenness. While degree centrality is usually a measure to characterize the importance of a node in a network, this is true only if immediate neighbors are the only ones determining the properties of a node. In contrast, betweenness here indicates how important the node is within the context of the entire connected component, and high betweenness lncRNAs, such as MALAT1, but also NEAT1 in the same component, might represent “essential” genes with a global effect in the leukemia gene regulatory network by orchestrating the regulation of distant gene clusters brought close to each other at chromatin level. While the physical interaction of lncRNA MALAT1 with the SIPA1 leukemia oncogene has been experimentally validated [38] and recapitulated in out network, MALAT1 is linked to other crucial oncogenes in the two linked modules. Among others we find EHBP1L1 and FOSL1, Rab effector proteins known to regulate cell proliferation, SYVN1, which has been shown to sequestrate the tumor suppressor p53 in the cytoplasm and thereby negatively regulates apoptosis [39], and the vacuolar protein VPS51 involved in endocytic recycling.

### Module structure and putative lncRNA functions in chromosome 17’s biggest component

Finally, we briefly discuss the clustering results of the biggest connected component of chromosome 17, obtained with *α* = 100 and *θ* = 0.7 (Fig. 5c), which shows a nice modularity, resulting in ten prominent modules, and it is particularly interesting because it contains several lncRNAs with very high degree/betweenness, so potential core players in the leukemia regulatory network, but of unknown function. The analysis results point, among others, to co-regulation of lncNRA LINC00910 and LINC00854 of unknown function, but both exhibiting very high degree and betweenness in their connected component, and clustered in the same regulatory module (cluster 8). Cases of lncRNA-lncRNA synergic interactions linking regulatory modules have been already observed in other studies [40]. LINC00910 is directly connected and putatively activating the DHX8 gene, shown to be required for correct cell division and genome stability in general [41]. In addition, it seems to connect several CTCF DNA binding elements (yellow nodes) and transcribed enhancers (green nodes), numerous in the LINC00910 module, to well-known tumor suppressor genes such as NBR1 and BRCA1, and another module (cluster 4) containing several known oncogenes such as ICT1, small ubiquitin-like modifier SUMO2, nuclear pore complex protein NUP85 and hormone-inducible gene HN1 involved in cancer progression and metastasis. Although the exact functions of LINC00910 and its partner LINC00854 remain to be determined, we can hypothesize that they are implicated in chromatin looping in complex with CTCF and other enhancer elements to activate or insulate the transcriptional activity of several cancer-related genes. Interesting is also the module containing the lncRNA RP11-400F19.8 of unknown function (cluster 6). This lncRNA connects the STAT transcription factors STAT3, STAT5A and STAT5B to several loci of genes which act as regulators of apoptosis and/or cell proliferation, such as PSMC3IP [42], VPS25 and MLX [43]. STAT factors are known to be important regulators of the Immune Response in a tumor environment in either promoting or inhibiting cancer, therefore here we hypothesize an important role of RP11-400F19.8 as activator of cancer genes, and a more general function as STAT-mediated regulator of the cancer immune response.

## Discussion

LncRNAs play key regulatory roles in a wide range of processes, and a small number of them has been shown to operate in the nucleus and influence transcriptional regulation of neighboring or distal genes. To which extent cell-type specific 3D chromatin organization and other DNA regulatory elements contribute to lncRNA-mediated gene regulation has been poorly investigated. In addition, functional annotation for most of the annotated lncRNAs, as well as their role in gene regulatory networks remains elusive. Based on the fact that transcripts sharing common expression patterns should largely share similar biological pathways, a number of different studies have used the ‘guilt by association’ approach to functionally annotated lncRNAs based on expression similarities with protein-coding genes of known function.

Here we comprehensively map ChIA-PET chromatin contacts mediated by PolII in the K562 cell line to lncRNAs, genes and other DNA regulatory elements, and propose a multi-step approach to analyze lncRNA regulatory function using graph analysis techniques. We focused on lncRNAs of high degree or betweenness centrality, as degree hubs are potentially at the heart of large regulatory modules, while nodes with high betweenness centrality are well-suited for mediating interactions within and between modules.

This is also the first study which attempts to identify lncRNA-mediated transcriptional regulatory modules by means of fuzzy clustering analysis on the chromatin network, and provides a first link between transcriptional reg-ulation and lncRNA association/functions at chromatin level. Although alternative choices exist for the module search, we decided on the MSM clustering since it does not impose an often artificial full partition of the network into modules. Instead, MSM outputs a fuzzy clustering which allows more flexible interpretation of lncRNA regulation. Also, the *θ* and *α* parameters of our clustering method, allow us to analyze nested sequences of node sets associated with modules and therefore define more or less relaxed or interconnected modules. These parameters are chosen to maximize the co-expression of genes and lncRNAs inside the same module.

Integration of co-expression and chromatin data gives a more direct readout of co-regulation or mutual regulation involving lncRNAs, by identifying direct lncRNA targets and the regulatory modules they belong to. Our analysis revealed both high-level properties of lncRNAs in connected components, such as a significant association between high degree and enhancer functions or disease association of lncRNAs, as well as detailed mechanisms of function inside individual modules, like the ones discussed for the wellknown lncRNAs MALAT1 and FALEC. In addition, our results show the potential of identifying novel functionally important lncRNAs, like the high degree lncRNA LINC00910 on chromsome 17 and its potential implication in chromatin looping via CTCF binding.

Incorporation of other Transcription Factor Binding Sites in the network annotation will in the future reveal even more details about co-factors involved in lncRNA-mediated transcriptional regulation. This will enable a better interpretation of individual modules or links between them. In addition, here we have shown some significant results for the PolII ChIA-PET network in the K562 cell line, and for the biggest connected component of each chromosome. However, our procedure is general enough to be applied to the chromatin interaction network mediated by other factors, for example CTCF, to other cell lines/tissues, and the module analysis repeated systematically for any other connected component. The module analysis reported in Additional file 2 provides already a starting point to investigate new potential lncRNA regulatory functions for dozens of annotated lncRNAs.

Although the modular lncRNA regulatory code remains to be tested, studying the modular regulation of lncRNAs, and investigating their connections to genes and other regulatory elements are important steps towards further definition of lncRNA functions on a system-wide level. The investigation of modules related to lncRNAs whose functionality is not yet known can suggest new targets and the regulatory components involved in regulation. Therefore, we propose that our functional annotation scheme can be applied to thousands of lncRNAs in a tissue-specifc manner.

## Conclusion

In this study we demonstrate that the integration of 3D chromatin interaction and co-expression analysis provides a powerful network analysis approach for *in silico* functional analysis of both known and novel lncRNAs involved in transcriptional regulation. The results presented here, in particular the detected regulatory modules on the ChIA-PET interaction network, are an important resource for further biological research.

## Declarations

### Acknowledgments

The authors kindly acknowledge Leonie Chiara Martens and Martin Vingron for insightful discussions.

## Funding

This study is supported by the DFG Grant MA 4454/3-1.

## Availability of data and materials

The datasets used and/or networks analysed during the current study are available from the corresponding author on request. Graph analysis has been performed using standard software and algorithms as cited in the text as well as MSM clustering, the pseudocode of which is publicly available via the cited publications.

## Competing interests

The authors declare that they have no competing interests.

## Author’s contributions

Conceived the idea: AM. Designed the study: DT, NC, HS, AM. Developed the MSM clustering algorithm: NC. Implemented the methodology, performed the statistical modeling and analysis: DT. Contributed to the analysis of the results: RP, HS. Wrote the paper: DT, NC, HS, AM. All authors read and approved the final manuscript.

## Additional Files

### Additional file 1

This pdf includes supplementary tables and figures referred to in the main text. This includes a figure with two small connected components with interesting lncRNAs as well as tables with network properties of lncRNAs in the biggest connected components of chromosomes 1, 11 and 17.

### Additional file 2

In this excel file we provide the results from clustering analysis of the biggest connected component of each chromosome, in order to assist future experimental studies. For each component, we report the results from those values of *α* and *θ* yielding the best partition according to the MI ratio criteria. We report the clustering parameters, the resulting MI ratio, the number of obtained modules per component, the number of lncRNAs, protein-coding gene, as well as the overall number of nodes for each component. Note when the genomic coordinates of a gene and a lncRNA overlap, both the gene and the lncRNA name are reported for the same node.

